# Dynamic Weight Signatures as a Novel Biomarker of Motor Dysfunction in Mouse Models

**DOI:** 10.64898/2026.11.23.768678

**Authors:** Amy Papaneri, Charles Tyler Shoemaker, Sukanya Saha, Shreya Ghosh, Guohong Cui

**Affiliations:** In Vivo Neurobiology Group, Neurobiology Laboratory, Division of Intramural Research, National Institute of Environmental Health Sciences, 111 TW Alexander Dr., Research Triangle Park, NC 27709, USA; University of North Carolina at Chapel Hill, Chapel Hill, NC 27599, USA

## Abstract

Motor dysfunction is typically assessed through clinical examination and manual scoring, with gait analysis and accelerometry providing objective measures of subtle abnormalities. The observable 4-6 Hz resting tremor of Parkinson’s disease highlights the importance of capturing motor fluctuations at relevant timescales. Using a high-precision industrial weighing module, we tested whether high-frequency weight recordings of freely moving mice could provide a simple, noninvasive means of detecting disease-associated motor abnormalities. Aging MitoPark mice spend more time in non-ambulatory states and exhibit less stable locomotion than healthy controls; however, bradykinesia and shaking can be mitigated by intraperitoneal levodopa. Here we demonstrate that irregular motor activity and tremor-like movements are quantifiable via weight frequency analyses, which permit detection of early symptoms and post-levodopa motor control restoration. This non-invasive approach is less data-intensive than video-based analyses, adaptable for home-cage monitoring, and potentially translatable for clinical or home use for detection of subclinical motor dysfunction.

## Introduction

Parkinson’s Disease (PD) is the second most common neurodegenerative disease affecting humans, presenting primarily as a movement disorder with other symptomatology commonly impacting cognition, sleep, digestion, and temperament. Parkinsonism is the main clinical motor manifestation of PD and is defined by bradykinesia in combination with rigidity and/or resting tremor. Additional symptoms of postural instability and gait impairment tend to emerge with disease progression [1],[2]. The related motor dysfunction reduces quality of life in PD patients, where parkinsonism leads to falls and related injuries that perpetuate the progressive health decline [3, 4].

First described by James Parkinson in 1817 as the “Shaking Palsy,” PD is often associated with a characteristic resting tremor typically occurring at a frequency of 4–6 Hz and initially affecting one hand [5–7]. Although not always present in PD, tremor is one of the most recognizable clinical features and, therefore, is often used as a measure for detecting disease amelioration. This resting tremor is typically observed only after significant dopaminergic neuron loss in the substantia nigra pars compacta (SNc), a hallmark pathological feature of PD [8]. Knowing this, earlier detection of parkinsonism would be beneficial for implementing measures of intervention to stop the progressive loss of dopaminergic (DA) neurons.

While conventional mouse behavioral assays, such as open field, rotarod, and gait analysis, measure locomotion and motor coordination, they do not readily enable investigation of subtle, high-frequency tremors. Accelerometry, alone or combined with electromyography (EMG), can provide visualization of different tremor classes [9, 10]; however, the data collection approaches require wearable devices and, when EMG is used, placement of multiple surface electrodes. Current strategies have advanced to include actometry-style mechanical tremor-detection systems [11–13] and body-worn sensors for monitoring PD patient symptoms remotely [14, 15]. Although tremor patterns at specific frequencies can provide clinically relevant indicators of disease, such as the 3-10Hz characteristic frequency range for non-walking tremor events in human studies [16, 17], cost-effective technologies suitable for widespread diagnostic and monitoring applications remain lacking.

Studies using mouse models of PD, as well as other motor disease models, have facilitated exploration of various diagnostic and treatment methods [18]. Considering the need for scalable, cost-efficient diagnostics and tremor monitoring methods, we sought to investigate how tiny changes in vertical force measurements of these behaving mice may reveal signs of motor dysfunction prior to the onset of clinical symptoms. We hypothesized that patterns of weight signatures reflecting motor dysfunction in these mouse models, potentially but not necessarily related to tremors, could be detectable using ultra-fast acquisition of high-sensitivity dynamic weight data. We tested the Mettler Toledo High Precision Weighing Module SPE404-C for its capability in rapid and sensitive acquisition of weight (i.e.: vertical force) data from freely moving mice. By analyzing these weight data recordings for signatures of healthy versus abnormal motor behavior, clinical diagnostic evidence of neurodegeneration may be feasible at an earlier age than via currently existing methods.

The progressively degenerative genetic mouse model of PD, MitoPark, exhibits DA cell loss in the SNc beginning at age 12 weeks with approximately 40% tyrosine hydroxylase (TH) cell loss in that brain region compared to control mice. By 33 weeks of age, MitoPark mice have lost approximately 80% of TH-expressing neurons in the SNc and displays impaired gait and bradykinesia [19]. Therefore, this model is practical for testing early detection methods of motor dysfunction as related to neurodegeneration. Additionally, our novel mouse strain cross for global cholinergic neuron disease, ChAT-tm1Cre/Tfam, exhibits rapid health decline involving gait abnormality and shaking from 13 weeks of age, with approximately 50% cholinergic neuron loss and debilitation necessitating euthanasia around 16 weeks of age (Saha, S. et al, manuscript in preparation).

Here we introduce this novel method for quantifying motor deficit in mouse models of disease by analyzing weight signatures recorded by an ultra-fast, high-precision weighing module. This non-invasive assessment for early detection of subclinical motor dysfunction has the potential to be developed for diagnosis and monitoring of degenerative motor diseases in veterinary settings and could be translatable to the human clinic.

## Results

### Ultra-fast, high-precision weight data aligns with behavioral video recordings

A Mettler-Toledo SPE404-C Weigh Module was used to record streaming weight data from MitoPark mice placed in a beaker on the platform (Fig1A). Ten minutes of recording were acquired at a rate of 122Hz. Video recordings were obtained using either a Teledyne FLIR Grasshopper 3 camera (FLIR-G3) or the Intel Real Sense D435i depth camera (D435i). Video provided behavioral context whereby movements could be aligned to weight signatures in order to validate the method.

**Figure 1.**
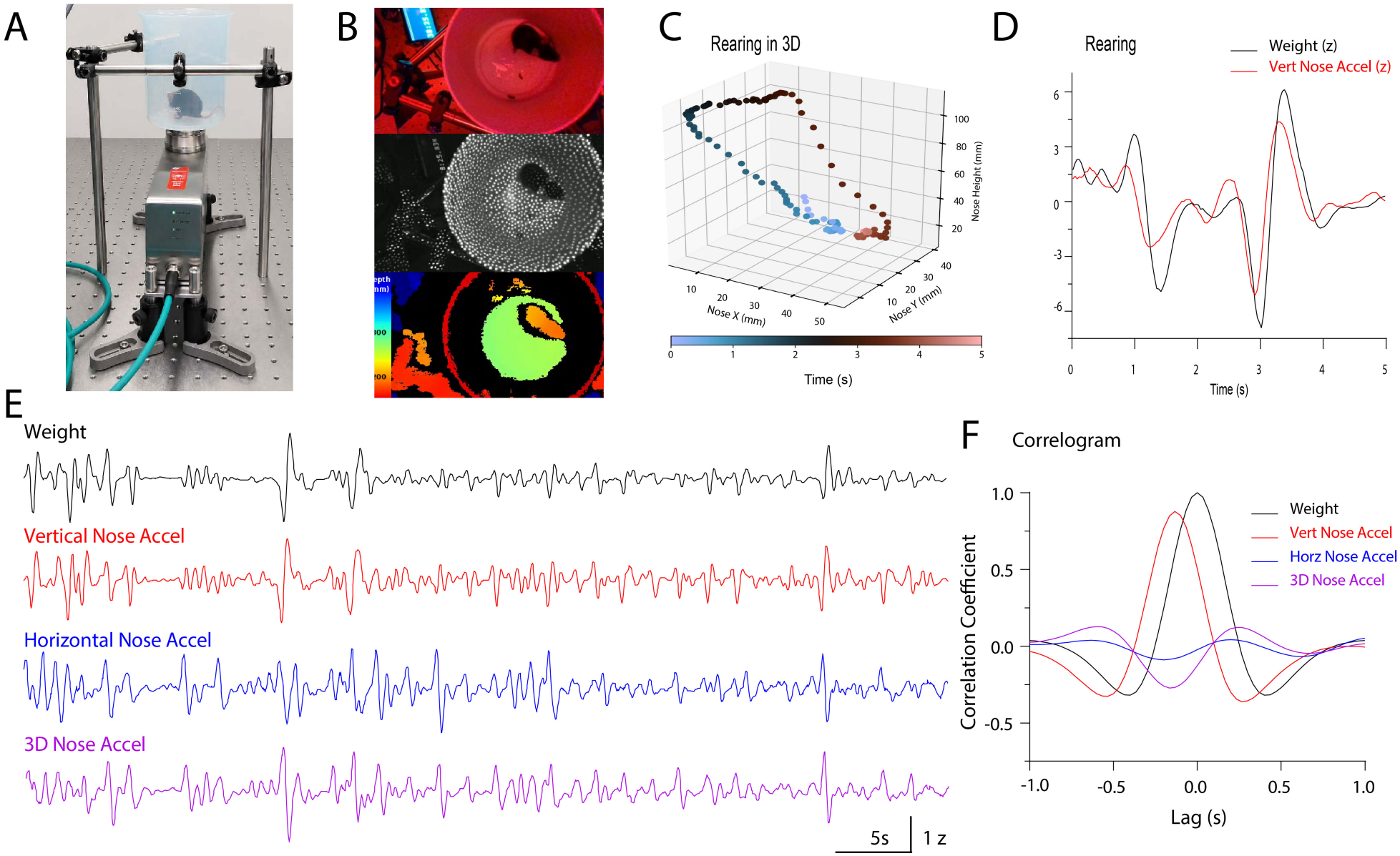
Hardware and alignment of high-precision weight data with movement. A) Image of Mettler Toledo SPE404-C high-precision weigh module setup. B) Overhead view from depth camera; top = visible light mode, middle = infrared dot projection, bottom = heatmap based on depth. C) 3D example of rearing over 5 sec of depth camera recording. D) Alignment of weight and vertical nose acceleration for the same rearing example as in C. E) Trace alignments for Z-scored weight, vertical nose acceleration (VNA), horizontal nose acceleration (HNA), and 3D nose acceleration (which is VNA+HNA) over 1min. F) Weight auto-correlogram with cross-correlations plotted for VNA, HNA and 3D nose acceleration from 10min recording.

Using the FLIR-G3 video, behavioral bouts could be distinguished broadly as periods of action versus inaction, whereas detection of specific behaviors, including grooming, forward or backward movement, rearing, resting, and stillness, was less consistent. After extensive effort in model training and optimization with limited improvement, we transitioned to the D435i for synchronized video recordings containing frame-by-frame position data.

The D435i depth camera permitted the acquisition of infrared depth recordings at 30 frames per second (Fig1B). Time-matched recordings of weight and behavior were subsequently analyzed using ChatGPT through a defined set of prompts (to be cited upon publication) with the resulting classifications compared against synchronized video observations. Mouse position was determined from the infrared video using a frame-by-frame image-processing procedure described in the Methods. We were able to effectively synchronize with our weight data to validate the performance of typical behaviors such as rearing (Fig1C and D). This example of rearing over five seconds of recording depicts how the weight dynamics during rearing closely correlates with vertical nose acceleration. Weight data was aligned temporally with mouse nose position (Fig1E). Freezing behavior also closely correlated with vertical nose acceleration, whereas horizontal nose acceleration and 3D nose acceleration were found to be inversely correlated in these instances Fig1F).

### Weight signature method detects restorative effect of L-DOPA in treated MitoPark mice

To test our weight signature method for motor dysfunction detection, we used the progressively degenerative genetic mouse model of PD, MitoPark (MP). By 34 weeks of age, MP mice had lost approximately 80% of TH-expressing neurons in the ventral tegmental area and substantia nigra pars compacta (VTA/SNc) as well as striatum (Fig.2A). Weight data traces were distinct for 33-to 34-week-old MP versus Littermate Control (LC) mice. Behaviors such as rearing were distinguishable, as noted in representative traces (Fig.2B).

Comparison of weight traces between genotypes allowed visualization of overall activity, which showed the reduced movement for 34w MP compared to LC and the restoration of greater amplitude activities following L-DOPA treatment (Fig2C). This change in overall activity level is quantifiable by evaluating the Coefficient of Variation (CV) and comparing pre- and post-LDOPA CV percentages. Weight recordings confirmed a significant difference between the CV for pre-LDOPA MP and LC mice, whereas CV at 30min post-LDOPA for MP was restored to that of LC (Fig.2D). Representative heatmaps of power spectral density (PSD) were generated in 10 sec bins for each 10 min recording to visualize the distribution of spectral power across frequencies throughout the session. Frequencies of 5-15Hz are most evident for LC mice, whereas MP mice exhibit a relative increase in spectral power between 15-22Hz (Fig.2E). The shortened timescale seen in Fig 2E is a result of cleaning the weight data and trimming to equal length (see Methods).

**Figure 2.**
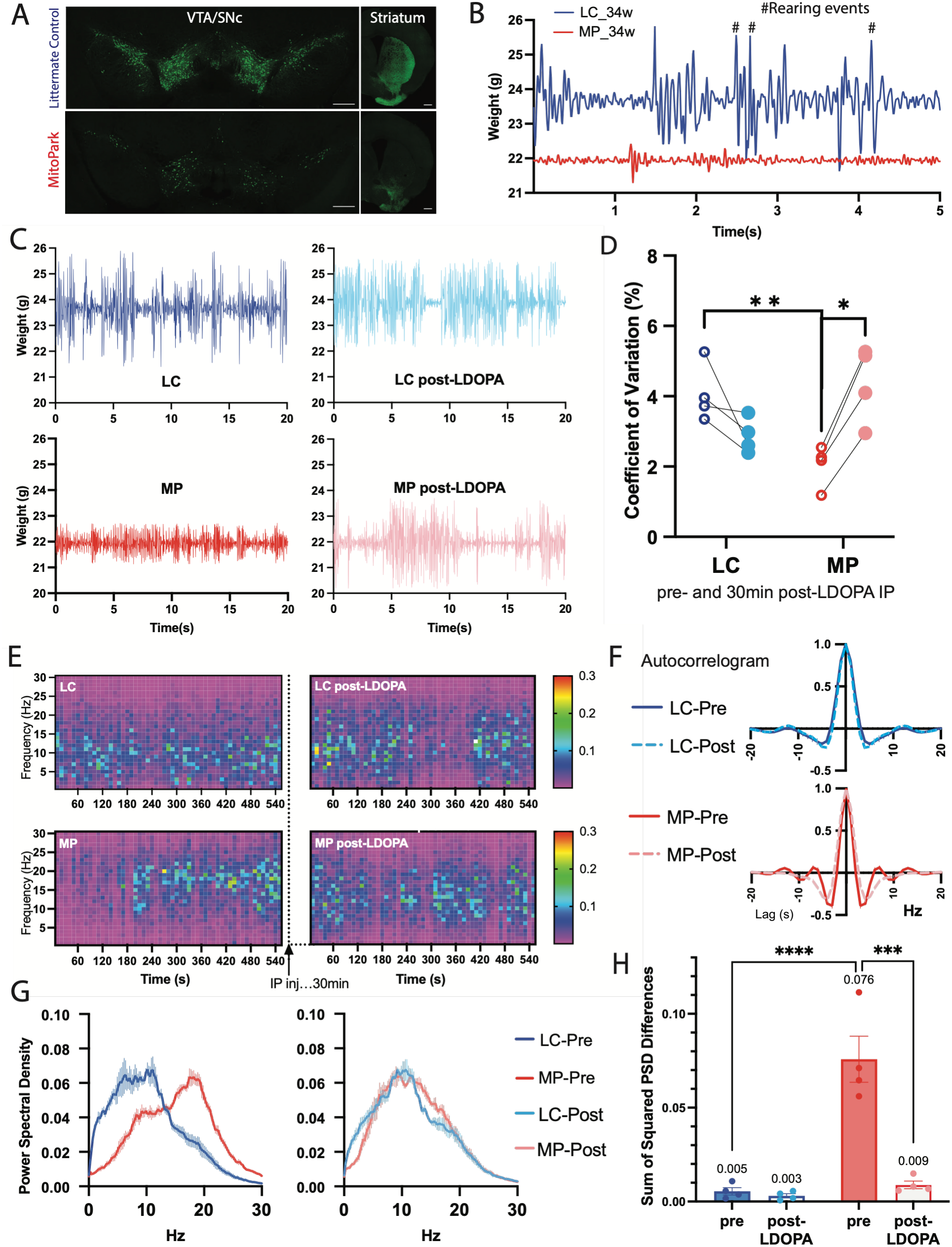
LDOPA treatment restores motor function in MP group based on ultra-fast, high-precision weight data for DAT-Cre/Tfam mice. For 33-34-week-old female Littermate Control (LC) and MitoPark (MP) mice, A) Representative confocal images of ventral tegmental area (VTA)/substantia nigra pars compacta (SNc) and striatum stained with anti-Tyrosine Hydroxylase (TH) primary antibody and Alexa488 secondary antibody. Scale bars, 200um for VTA/SNc; 50um for Striatum, and B) representative 25 sec of raw weight recording. Analyses of 10min weight recording data from LC or MP mice, pre- and post-LDOPA IP administration, N=4/group. LC = blue; MP = red; LC-post-LDOPA=light blue; MP post-LDOPA=pink. C) Representative 20 sec; D) Coefficient of Variation; E) Heatmaps of PSD frequencies; F) Autocorrelation across ±20sec lags; G) PSD frequency distribution from 0 to 30Hz; H) Sum of the Squared PSD Differences in G. Data are presented as mean ± SEM. Group differences were assessed using a two-tailed unpaired t test. P-values are reported where relevant. Groups: LC = blue; LC post-LDOPA= light blue; MP = red; MP post-LDOPA = pink. Significance values: *=p<0.05; **=p<0.01; ***=p<0.001; ****=p<0.0001.

Auto-correlograms comparing pre- and post-L-DOPA weight signals reveal a steeper decline and non-conforming decorrelations for MP mice, indicative of erratic behavior, particularly evident at lags 4 and 7. The trough at lag 4 followed by zero-crossing near 7 are consistent with an oscillatory component in the 13-15Hz range. Following L-DOPA administration, these peaks smoothed to align closely with LC auto-correlograms, providing confirmatory evidence of the restoration of normal behavioral patterns (Fig.2F).

Power spectral density (PSD) graphs of 33-34w weight-log data from 10 min recording sessions also provide a clear illustration of the shift in frequency domain that occurs in aging MP mice. Whereas spectral power for LC is predominantly concentrated between 1 and 12Hz, MP spectral power is reduced in this range, and a second peak at 15-20Hz is evident (Fig. 2G). By dosing IP with L-DOPA and recording another 10 min session at 30min post injection, PSD for MP shifts to closely resemble that of LC. The transformation in PSD can be quantified by summing the squares of the differences between the LC-Pre value and the values for LC-Post, MP-Pre and MP-Post, where “Post” is the 10 min recording started at 30min post-IP injection of L-DOPA. These results indicate that L-DOPA restored the overall PSD profile of 34-week-old MP mice to a pattern comparable to that observed in LC mice (Fig.2H).

### Novel method detects motor deficit early for the PD mouse model MitoPark

Previous studies have not yet successfully quantified dysfunction in MP mice any earlier than 12w [2, 20, 21]. To evaluate whether the weight data collected by the SPE404-C could reveal disease condition at an earlier age in MP mice, 10 min weight recording sessions were performed on a cohort of DATTf mice at ages 8w, 10w, and 12w.

Calculation of the coefficient of variation (CV) confirmed that overall diminished activity is detectable for MP at 12w via this novel method, while significant differences in overall force measurements are not evident at 8w or 10w (Fig.3A). Power Spectral Density graphs were nearly identical for LC and MP at 8w and 10w. However, the MP curve shifts rightward at 12w, indicating increases in activity at higher frequencies and revealing the first emergence of the 15-20Hz peak (Fig.3B). Sum of squared PSD differences provide further evidence that significant change in MP frequency domain is quantifiable at 12w (Fig.3C). Again, autocorrelation results at 8w and 10w are nearly identical between LC and MP, but significant differences emerge at 12w (Fig.3D). The main decorrelation is evident at lag 4 for 12w MitoPark mice (Fig.3E). Although our evaluations have not yet identified a statistically significant difference in MP mice at a timepoint earlier than 12w, detection at 10 or 11 weeks could still be plausible via this method with a larger sample size and analysis focused on a targeted frequency range.

**Figure 3.**
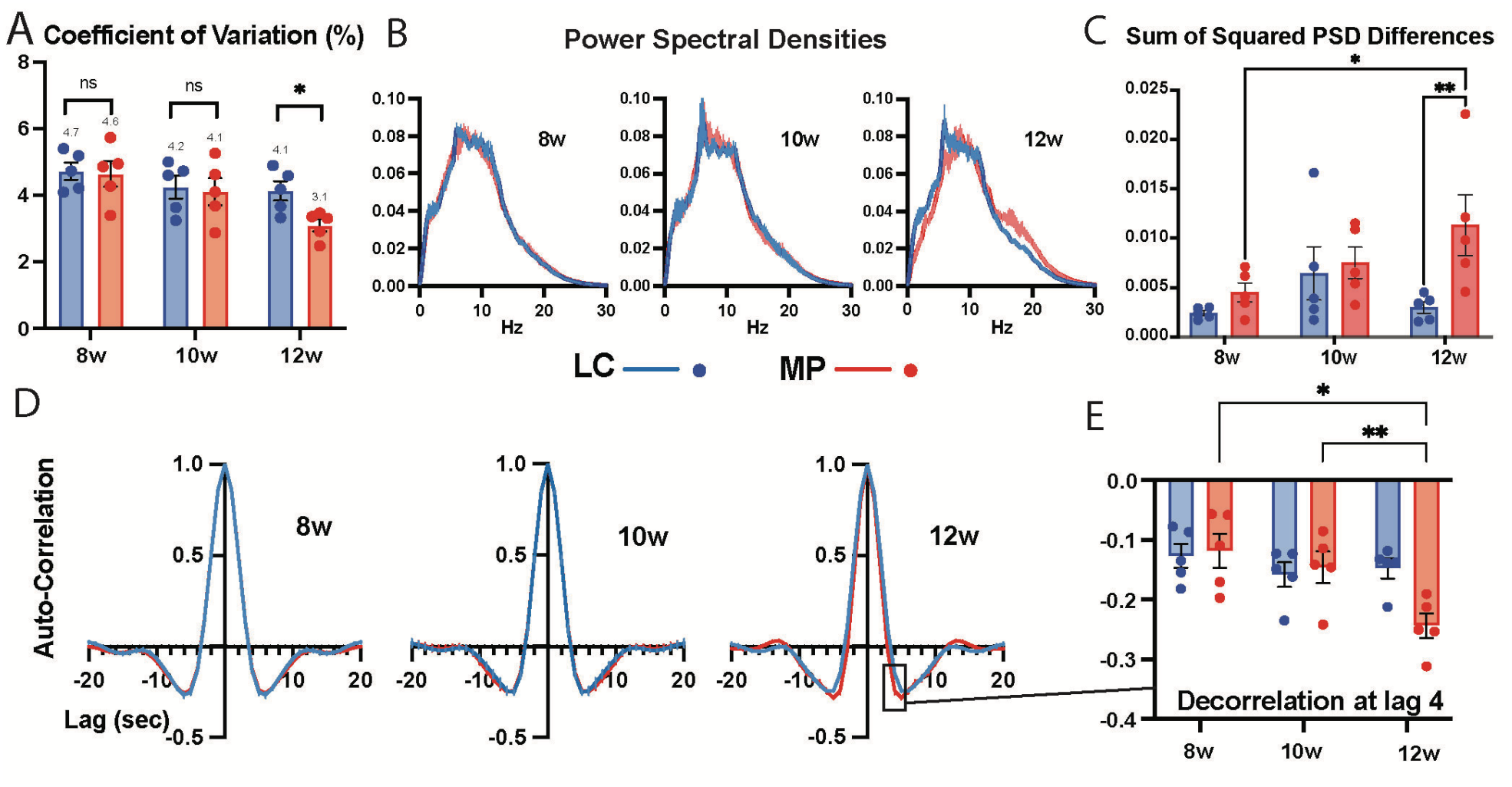
Multiple analyses of 10min recording data from LC and MP groups at ages 8w, 10w, and 12w; N=5/group. A) Coefficient of Variation; B) Power Spectral Density charts; C) Sum of squared PSD differences; D) Auto-correlograms; E) Selection of significant decorrelation emerging for MP at 12w. Bar graphs are mean values with points representing individual mice; error bars are +/-SEM. LC = blue; MP = red. Significance values: *=p<0.05; **=p<0.01.

### Control results are confirmed as comparable via reproducibility in WT background strain

Optimally, the use of this method with varied mouse strains would produce distinguishable, strain-specific weight signatures, with mice of the same genetic background being expected to exhibit greater similarities in their signatures. Therefore, in order to validate the utility of this novel method for screening across multiple mouse lines, DATTf mice were compared to their background strain, C57BL/6J, at 25w of age. A small cohort of 4 or 5 mice group (at least two of each sex) were screened to obtain preliminary data. Despite the small N, this method produced quantifiable and statistically significant results for overall activity level (Fig.4A), with C57 mice exhibiting an average CV of 4%, while DATTf-LC were more active at 5.8% CV and MP were less active at 2.8% CV. Autocorrelation for each strain was plotted for these 25w old mice, and the C57 auto-correlogram aligned nicely with the DATTf-LC result, whereas the data for MP mice produced a steep decline with a significant trough at lag 4 and a peak at 7 (Fig.4B). This ACF pattern was very similar to that for 33-34w old MP mice and may represent a behavioral signature, although confirmation is still pending further analysis. Graphing the PSD again illustrates the characteristic dual peaks for MP mice at 25w, whereas LC and C57 are highly aligned, and the majority of their spectral power is between 1 and 12Hz (Fig.4C). Quantification of the PSD differences was calculated by summing the squared differences between C57 group values and those of either LC or MP mice (Fig.4D). The overall change in PSD for 25w MP mice compared to controls is on the same order of magnitude as the comparison at 33-34w, while the change at the 12w timepoint is an order of magnitude smaller yet still significantly different from LC. This shifting of power for MP mice from lower to higher frequency, which began to emerge at 12w of age (Fig.3B), must be investigated to determine how it relates to or signifies disease-condition.

### Unique weight signature results for a mouse model of cholinergic disease, ChAT-Cre/Tfam

To confirm that this method can serve to potentially diagnose mouse models of motor disease, we tested a novel mouse cross that was developed based on the same cre-dependent mitochondrial transcription factor A deletion strategy as MitoPark mice. ChAT-tm1cre/Tfam homozygous mice lose cholinergic neurons globally, and knockout mice for this strain were termed Cholinergic Neuron Disease (CND) mice. This depletion of cholinergic cells causes rapid progression of motor dysfunction between 13-16w of age, as evident from ChAT-stained brain slices from 14w CND and LC mice (Fig.5A). CND and LC mice were weighed on the SPE404-C for 10 min recording sessions. A representative 5 sec trace for the same mice used in Fig.5A is shown in Fig.5B, showing the sporadic nature of the activity of CND mice with disease progression. Unlike MitoPark mice, differences in overall activity level of ChAT-tm1cre/Tfam mice based on coefficient of variation of 10 min weight recording data are not evident for CND mice versus age-matched littermate control mice (Fig.5C). While overall activity level was unchanged, autocorrelation analysis revealed differences in detected vertical force for this strain, especially at lag of 4 and 7. Despite group sizes being too small to analyze for sex differences, a trend indicated that males have higher CV values than females for this strain (data not shown), yet values were not significantly different as a function of time either for males or for females.

Other frequencies of interest, along with the highlighted lag 4 discrepancy, have yet to be assessed for the behavioral implications. For example, abnormal peaks at 7 and 14 lag are also evident for MitoPark mice at ages at 25w+ as seen in Figures 2 and 4. Aside from changes in gross motor behavior, changes in sniffing and whisking behavior should be evaluated as a potential cause for these discrepancies, given the biologically relevant timescales of ~10 and 20ms, respectively.

**Figure 4.**
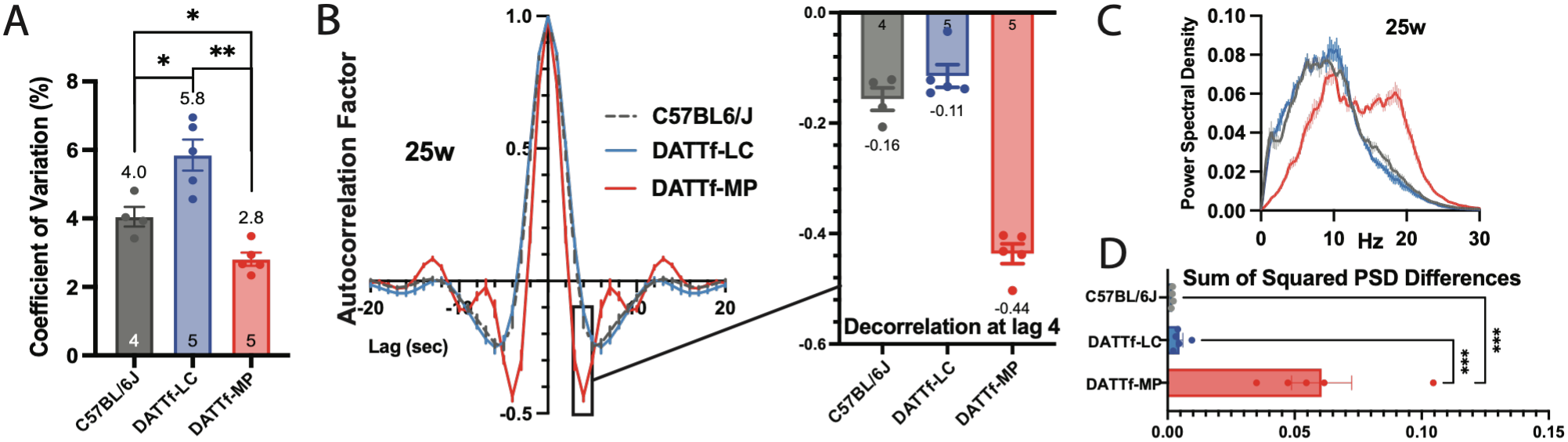
DATTf LC and MP mice compared to C57BL/6J at 25w of age. A) CV; B) ACF; C) PSD; D) Sum of Squared PSD Differences. C57 = gray; LC = blue; MP = red. Bar graphs are mean values with points representing individual mice; error bars are +/-SEM. N values are inside the bars. Significance values: *=p<0.05; **=p<0.01; ***=p<0.001

**Figure 5.**
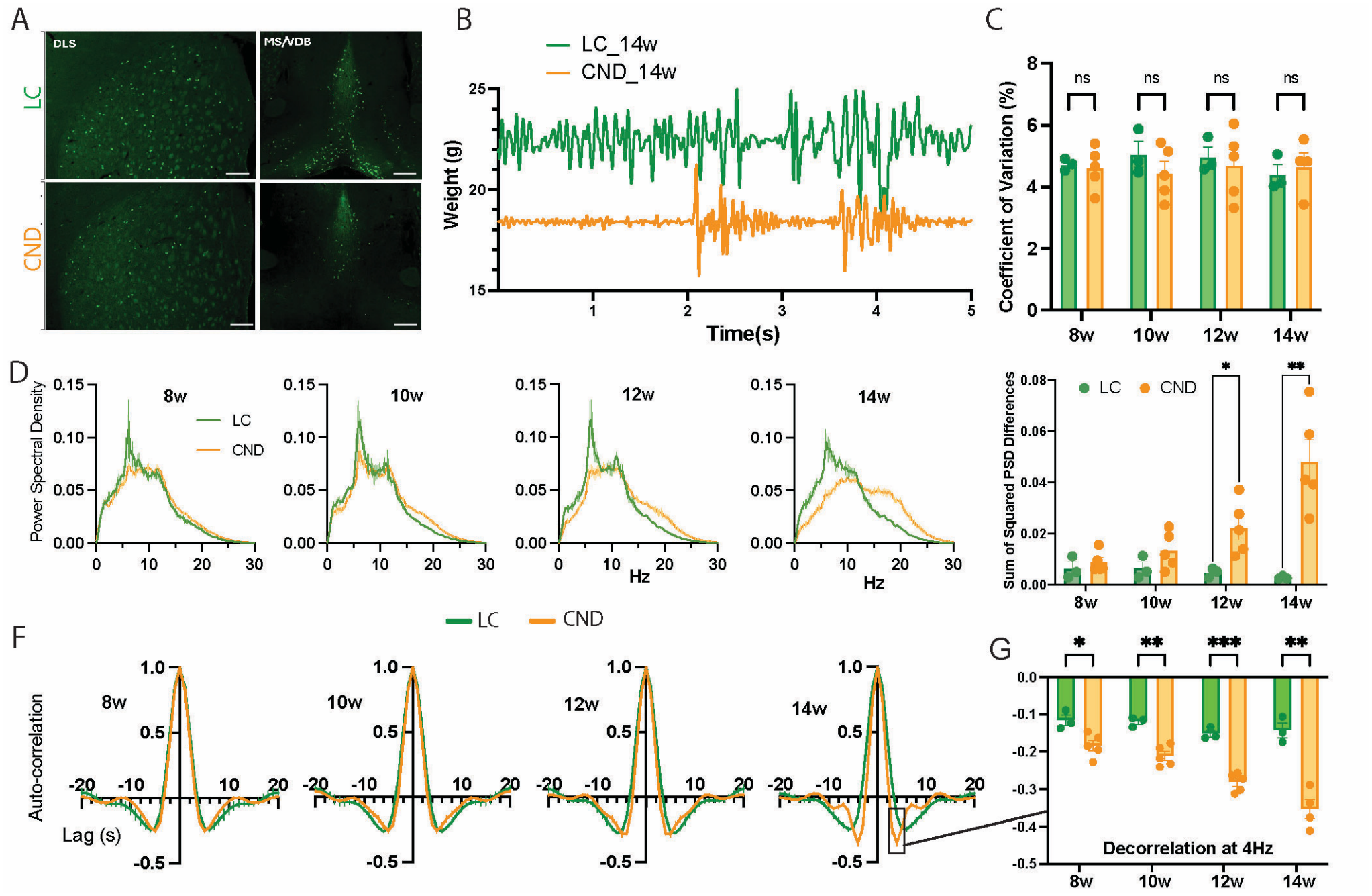
Results for a cholinergic-knockout mouse model. Representative 20x confocal images of dorsal-lateral striatum (DLS) and medial septum/ventral diagonal band of Broca (MS/VDB) from 14-week-old ChAT-Cre/Tfam mice, Littermate Control (LC) and Cholinergic Neuron Disease (CND), stained with rabbit anti-ChAT primary antibody and anti-rabbit Alexa488 secondary antibody. Scale bars: 50um for DLS, 200um for MS/VBD. B) Raw weight values from 14-week-old female ChAT-cre/Tfam mice, representative five seconds of recording. C) Coefficient of Variation, changes in overall force measurements between LC and CND. D) Power Spectral Density (PSD) comparisons. E) Mean squared differences in PSD between LC and CND at ages 8, 10, 12 and 14 weeks old. F) Auto-correlograms (AC) for LC and CND mice at ages eight, ten, twelve and fourteen weeks. G) Highlighting the decorrelation at 4Hz for CND mice at all ages tested. LC = green; N=3. MP = orange; N=5, except at 14w where N=4. Bar graphs are mean values with points representing individual mice; error bars are +/-SEM. Significance values: *=p<0.05; **=p<0.01; ***=p<0.001.

Supplemental data using Mettler Toledo PBK989APW-A3 platform scale (weight signatures, CV, AC, PSD) will be added in a subsequent version of this manuscript.

## Discussion

Parkinson’s disease (PD) is predominantly idiopathic or sporadic, with disease risk thought to reflect interactions among genetic susceptibility, environmental exposures, and aging-related processes that may begin years or decades before the emergence of clinically apparent motor symptoms [22, 23]. Although PD primarily affects older adults, approximately 5% of cases in the United States are classified as early- or young-onset PD, some of which are associated with genetic, drug-related, or other disease-related factors [24, 25]. Recent epidemiological analyses further suggest that the global incidence and prevalence of early-onset PD may increase, emphasizing the need for approaches capable of identifying motor dysfunction at earlier stages and monitoring disease progression with minimal burden [26, 27]. Similar interactions among genetic susceptibility, environmental insults, neuronal dysfunction, and aging contribute to motor abnormalities in other neurodegenerative conditions, underscoring the broader value of sensitive and scalable approaches for quantitative motor phenotyping.

The present study demonstrates that high-frequency, high-precision measurements of dynamic body weight can provide clinically relevant information about motor function in mice. Using the Mettler-Toledo SPE404C weigh module, we captured mechanically transmitted signals generated by freely moving mice at high temporal resolution and sensitivity. These dynamic weight signatures distinguished behavioral states and revealed disease-associated differences in movement magnitude, temporal organization, and frequency content in mice. Importantly, for the MitoPark (MP) model, abnormalities were evident before or during the development of overt motor impairment and shifted toward control-like patterns following levodopa/carbidopa treatment. A mouse model of cholinergic dysfunction was also evaluated and produced quantifiable strain-specific results (ChAT-tm1Cre/Tfam; CND). Together, these observations suggest that high-frequency dynamic weight measurements can provide an objective, minimally invasive readout of motor phenotypes arising from distinct forms of neuronal dysfunction and pharmacological response, although additional work is needed to establish the specificity of individual signal features across disease models.

An important advantage of this approach is that substantial information about animal behavior can be extracted from a single, relatively simple mechanical signal. Conventional behavioral analysis often relies on video-based tracking systems that generate large datasets and may require extensive model training, annotation, storage, and computational processing. During method development, we evaluated video-based behavioral tracking approaches, including via DeepLabCut and EthoVision XT, but we found that reliable classification of several behaviors required considerable optimization. Synchronized video remained valuable for validating behavioral states identified from the weight traces; however, the present results suggest that, once characteristic weight signatures are established, many broad behavioral states can be distinguished without continuous reliance on computationally intensive video analysis. This reduction in data and processing requirements may facilitate higher-throughput studies and longer-duration monitoring.

The analytical workflow is also readily scalable. Raw high-frequency weight records can be batch processed using open-source programming languages such as R or Python, allowing automated extraction of behavioral, spectral, and temporal features and generation of summary statistics and visualizations. Thus, although acquisition produces a relatively large volume of raw data, the manual burden associated with data collection and downstream analysis can remain low once standardized processing pipelines are established. Automated analysis may be particularly valuable for longitudinal experiments in which repeated measurements are required across ages, disease stages, or treatment conditions.

The ability to extract frequency-domain information from dynamic weight signals may be particularly useful for investigating subtle motor abnormalities. In addition to the prominent disease-associated difference in auto-correlation identified at lag 4, MP mice exhibited additional spectral and auto-correlation oddities compared to LC, including abnormalities near 7 and 14 lag and the power spectral density peak in the 15–20 Hz range that emerged at 12w. The behavioral and physiological origins of these signals remain to be established. Accordingly, these signals are best interpreted as disease-associated oscillatory motor signatures rather than definitively identified as tremor-related phenotypes.

This method may also be useful for evaluating interventions that alter motor function. Exercise is increasingly incorporated into the management of PD and has been associated with improvements in mobility, balance, and other motor outcomes [28, 29]. Longitudinal dynamic weight measurements could therefore provide an objective means of assessing the effects of exercise or other behavioral interventions in MitoPark mice, including studies incorporating voluntary running-wheel activity. Similarly, the approach may be useful for quantifying acute and chronic effects of pharmacological agents on coordination, movement stability, and tremor-like activity.

Several limitations should be considered. First, the SPE404C system and associated acquisition hardware may be cost-prohibitive for some laboratories, particularly when multiple recording stations are required. We are therefore evaluating lower-cost implementations that preserve sufficient temporal resolution and sensitivity while substantially reducing equipment costs. Second, the present method measures forces transmitted through the weighing platform rather than movement of individual body segments. Consequently, the anatomical source of a particular oscillatory signal is difficult to distinguish from the weight trace alone.

Our findings extend previous work showing that movement-sensing platforms can provide quantitative measures of rodent activity and tremor-like movement [30–32]. However, many of these existing approaches require transfer of the animal to a dedicated recording apparatus, which may alter spontaneous behavior and limits the duration over which natural activity can be assessed. An important future direction is therefore integration of high-frequency weight sensing into the home-cage environment. Continuous or repeated home-cage measurements could reduce handling-related perturbations while providing longitudinal information about spontaneous movement, rest, tremor-like activity, and disease progression.

Despite these limitations, the simplicity and noninvasive nature of the approach create potential translational opportunities. Human movement monitoring increasingly relies on wearable accelerometers, inertial sensors, and other devices for assessment outside the clinic [33–35]. High-frequency force or weight sensing via brief platform interaction offers a strategy that could potentially obtain motor information without requiring a device to be worn continuously. If analogous disease-associated signatures can be detected in humans, high-resolution sensing incorporated into household or clinical scales could enable brief, repeated assessments of motor stability, tremor-like activity, medication response, or functional decline.

Overall, these findings demonstrate that high-frequency dynamic weight measurements can be extracted from mechanically transmitted signals using relatively simple acquisition and automated computational analysis, with potential as a novel biomarker enabling detection of disease-associated and treatment-responsive motor signatures. Further validation of the physiological origins and disease specificity of these signals, together with development of lower-cost and home-cage-compatible hardware, will determine the extent to which this approach can be used for longitudinal preclinical phenotyping and ultimately adapted for low-burden monitoring of human motor dysfunction.

## Materials and Methods

### Animals

All animal procedures were approved by the Institutional Animal Care and Use Committee of the National Institute of Environmental Health Sciences (NIEHS). Experiments were performed using male and female mice aged 2-9 months. Mice were singly housed after 8–10 weeks of age in a reversed 12:12h dark/light cycle room at 18–23 °C with 40–60% relative humidity with ad libitum access to standard chow and water.

MitoPark (MP) mice (DAT^*IREScre/+*^;Tfam^*flox/flox*^; also referred to as DATTf) were generated and bred in-house through a two-step crossing strategy between DAT^*IREScre/+*^ mice (The Jackson Laboratory, stock number 006660) and Tfam^*flox/flox*^ mice (The Jackson Laboratory, stock number 026123) as previously described [36]. Tfam^*flox/flox*^ littermates were used as negative controls.

Cholinergic Neuron Disease (CND) mice (ChAT^*tm1cre/+*^;Tfam^*flox/flox*^; also referred to as ChTf) were generated and bred in-house through a two-step crossing strategy between ChAT^*tm1cre/+*^ mice (The Jackson Laboratory, stock number 031661) and Tfam^*flox/flox*^ mice (The Jackson Laboratory, stock number 026123). Homozygous ChAT^*tm1cre/tm1cre*^ mice were crossed with homozygous Tfam^*flox/flox*^ mice to obtain ChAT^*tm1cre/+*^;Tfam^*flox/+*^ offspring. These heterozygous mice were then crossed with homozygous Tfam^*flox/flox*^ mice to obtain ChAT^*tm1cre/+*^;Tfam^*flox/flox*^ for use in experiments. Tfam^*flox/flox*^ littermates were used as negative controls.

C57BL6/J mice (C57; also referred to as WT) were bred in-house from breeding stock obtained from The Jackson Laboratory (stock number 000664).

### Instrumentation

The Mettler-Toledo SPE404-C Weigh Module was mounted to an air table using threaded connectors. The weigh module was connected to a SuperMicro workstation (SMW, 2015 model CSE846E1R900B) via ethernet cable and Power-over-Ethernet (PoE) injector.

Initially, video recordings of mouse behavior were obtained using a Teledyne FLIR Grasshopper 3 camera and synced to weight log data based on timestamp. Later, for synchronized video recording of 3D position, an Intel RealSense D435i depth camera was used along with the SMW clock.

### Weight recordings

APWLink software (METTLER TOLEDO, Greifensee, Switzerland) was used to detect configuration and adjust the settings of the weigh module to maximize the capability for recording the unstable weight. Initially, the weight value was tared using Normal Weighing Mode for stability. Then the mode was switched to Sensor Mode with settings of “Very Unstable” and “Limit 20”, which was the maximum cutoff, in order to acquire rapidly varying weight signals. A 1L polypropylene beaker (VitaLab) was centered on top of the weighing pan and tared. Holding the mouse by the tail, the mouse was lowered inside and placed on the bottom of the beaker. Yet Another Terminal (YAT) open-source software (SourceForge) was used to record the streaming weight data at the acquisition rate of 122Hz. After ten minutes of recording, the log was turned off, and the mouse was returned to its home cage. Mice receiving L-DOPA IP were injected with 4mg/kg levodopa (Sigma) in sterile saline + 15mg/kg carbidopa (Sigma), which was first dissolved in dimethyl sulfoxide (DMSO, Cell Signaling Technologies), for final DMSO concentration of 2% (mixture termed L-DOPA).

### Video Recordings

Video recordings were obtained using either a Teledyne FLIR Grasshopper 3 camera (FLIR-G3) or the Intel Real Sense D435i depth camera. FLIR-G3 video was evaluated manually or used for model training via DeepLabCut (DLC; RRID: SCR_021391) or EthoVision XT (Noldus Information Technology, Wageningen, The Netherlands; RRID: SCR_000441) [37–40]. Behavioral bouts could be distinguished broadly as periods of action versus inaction, whereas detection of specific behaviors, including grooming, forward or backward movement, rearing, resting, and stillness, was less consistent. After extensive effort in model training and optimization with limited improvement, we transitioned to the Intel RealSense Depth Camera D435i for synchronized video recordings. Time-matched recordings of weight and behavior were subsequently analyzed using ChatGPT through a defined set of prompts (to be cited upon publication) with the resulting classifications compared against synchronized video observations. These analyses supported the ability of dynamic weight traces to distinguish broad mouse behavioral states without requiring continuous reliance on data-intensive video recording, storage, and processing.

The Intel RealSense D435i depth camera was mounted facing downward approximately 27 cm above the base of the beaker. The camera was connected via USB 2.1 and controlled using Intel RealSense Viewer software. Images were acquired at 30 frames/s with a resolution of 480 × 270 pixels using the infrared (Y8) and depth (Z16) channels. The camera’s laser emitter and automatic exposure functions were enabled during acquisition. To synchronize the camera recordings with the weigh module, a secondary 5-inch monitor was positioned within the camera’s field of view and displayed the PC clock at millisecond resolution. The monitor had a refresh rate of 66 Hz, and the displayed time was advanced by 23 milliseconds (ms) to compensate for the estimated 1.5-cycle display refresh latency. For this video recording experiment, a matte white cylindrical container of approximately 1L volume was used, instead of the translucent VitaLab 1L beaker due to its mildly reflective surface, to facilitate reliable depth-camera imaging.

If video was being recorded simultaneously, the video recording was started first. The YAT log was then immediately turned on to begin saving the streamed weight data to a .log file. After ten minutes, the log was turned off, and the saved file was transferred to a permanent storage volume. The first computer-generated timestamp in the YAT log was used as t0.

Mouse position was determined from the infrared video using a frame-by-frame image-processing procedure. Each infrared frame was converted to a binary image using an intensity threshold of 35 on an 8-bit scale (0–255). The image was first restricted to the largest contiguous white region, after which all black regions except the second largest were removed to isolate the mouse from other features in the field of view. For each frame, the black pixel located most centrally relative to the remaining black pixels was identified as a central reference point. The black pixel located farthest from this central point was then identified as one endpoint of the animal, and the farthest black pixel extending in the opposite direction was identified as the second endpoint, with the three points required to form an angle >60°. These endpoints were used to estimate the nose and tail positions and thereby characterize the position and orientation of the mouse. Finally, the resulting trajectories were manually postprocessed to correct erroneous nose–tail inversions and abrupt, artifactual jumps in the estimated nose position.

### Immunofluorescence

Mice were anesthetized via intraperitoneal injection of pentobarbital at a dose of 100 mg/kg per NIEHS SOP. Animals were then transcardially perfused with 1x phosphate buffered saline (PBS), followed by 4% paraformaldehyde (PFA). Brains were carefully extracted and post-fixed in 4% PFA overnight at 4°C, then transferred to 30% sucrose solution at 4°C for cryoprotection and further histological processing. Next, the brains were coronally sectioned at a thickness of 35 μm using a vibratome (Microm HM 430). Sections were collected into a 24-well plate containing 0.5 ml PBS per well. To visualize tyrosine hydroxylase (TH), coronal brain sections were incubated overnight at 4°C on an orbital shaker with rabbit polyclonal anti-tyrosine hydroxylase (TH) primary antibody (Millipore AB152; 1:1000 dilution). The following day, sections were washed with PBS and incubated for 1 hour at room temperature with Alexa488-conjugated anti-rabbit secondary antibody (Life Technologies A-11034; 1:500 dilution), again on an orbital shaker. Sections were subsequently washed with PBS. Finally, brain sections were mounted onto microscope slides using hard-set mounting media (VectorLabs H-1500), covered with coverslips, and allowed to dry at room temperature for 15–30 minutes. Slides were then stored at 4°C until imaging. Images were acquired using a confocal microscope at 10x or 20x magnification and processed with Zen Blue software.

### Data Handling

Batch processing of raw .log files was performed using RStudio to generate comma-separated-value (.csv) files for downstream analysis. Once standardized, this workflow enabled large numbers of recordings to be processed with limited manual intervention and supported automated generation of summary statistics, spectral analyses, autocorrelation analyses, and graphical outputs. Files generated during batch processing were transferred to designated storage directories after completion to prevent accidental overwriting or unintended inclusion in subsequent batch summaries.

Using APWLink software to configure the device, the SPE404-C weighing module detected changes in mouse weight at a readability of 0.1mg, and a YAT log recorded the changes in weight at a rate of acquisition of 122 times per second (Hz). Recording sessions lasted ten minutes each, and the resulting weight data, 600s at 122Hz, were converted to CSV format using a custom R script executed in RStudio (Posit Software, PBC, Boston, MA, USA; RRID: SCR_000432). To describe the process briefly, the 73200+ data points per recording were cleaned to remove outliers, i.e.: negative values and values greater than ten standard deviations to account for rare instances when mice jumped, were briefly airborne (zero or negative values), and then landed, exerting greater force onto the scale than consistent with the behaviors being tracked for analysis in this study. Weight logs were then trimmed to the same length for all recordings in the subset analysis group for truthful comparison.

To compare relative weight patterns over time across animals while minimizing differences attributable to absolute body mass, weight signals were normalized except where reported as raw. Weight values were either mean-normalized or Z-scored depending on downstream processing use. Mean-normalized values were calculated as:

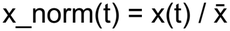

where x(t) represents the recorded weight at time t and 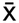 represents the mean value for that recording.

Mean-normalized data were used for Auto-Correlation graphs. Autocorrelation function (ACF) values were generated using an R script applied to the .csv weight-log data sampled at 122Hz. At this sampling rate, each lag corresponds to a time offset of approximately 8.2 ms (1/122 s). Recurring ACF features can be used to estimate periodicity and the corresponding signal frequency.

For power spectral density (PSD) analysis, signals were Z-score standardized according to:

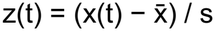

where 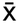 is the mean and s is the standard deviation of the recording.

Z-score standardization centered each signal at zero and scaled it to unit variance, thereby minimizing differences in overall signal amplitude and permitting comparison of how signal variation was distributed across frequencies. Accordingly, PSD analyses of Z-scored signals were interpreted as comparisons of spectral shape and relative frequency content rather than absolute movement power.

### Statistical Analysis

Investigators were blinded during data collection and analysis when feasible. GraphPad Prism software version 11 was used for graphing and statistical analyses of data. Welch’s unpaired t-tests were used for comparisons between two groups, whereas ANOVA was used for comparisons among more than two groups. Statistical significance was defined as α = 0.05, with Holm–Šidák correction for multiple comparisons where appropriate. Power spectral density (PSD) was estimated using a 128-sample Hann window with 50% overlap.

## Data and Code availability

Analysis codes used for the findings of this study will be made available at GitHub (https://github.com). Custom R [41] scripts have also been archived. Repository accession information and links will be added in a subsequent version of this manuscript.

## Acknowledgements

This research was supported by the Intramural Research Program of the NIH, National Institute of Environmental Health Sciences of the United States (ZIA-ES103310 to G.C.). The contributions of the NIH authors are considered Works of the United States Government. The findings and conclusions presented in this paper are those of the authors and do not necessarily reflect the views of the NIH or the U.S. Department of Health and Human Services.

We thank Mike Keith at Porter Scales, Raleigh, NC, and the technical experts at Mettler Toledo for their assistance with the equipment; Kelly Miller of the NIEHS Library for her assistance with literature searching; the NIEHS Neurobehavioral Core for their assistance with EthoVision XT; and the NIEHS Comparative Medicine Branch for their assistance with animal husbandry.

## Author Contributions

A.P.: Investigation, Methodology, Data curation, Formal analysis, Visualization, Writing – original draft, review & editing. C.T.S.: Investigation, Methodology, Data curation, Validation, Writing – review & editing. S.S.: Immunofluorescence and Confocal microscopy, Writing – review & editing. S.G.: Investigation, Data curation. G.C.: Conceptualization, Resources, Supervision, Writing – review & editing.

## Competing Interests

The authors have no competing interests.

## Notes

### Competing Interest Statement

The authors have declared no competing interest.

